# The phosphatase inhibitor Sds23 regulates cell division symmetry in fission yeast

**DOI:** 10.1101/632349

**Authors:** Katherine L. Schutt, James B. Moseley

## Abstract

Animal and fungal cells divide through the assembly, anchoring, and constriction of a contractile actomyosin ring (CAR) during cytokinesis. The timing and position of the CAR must be tightly controlled to prevent defects in cell division, but many of the underlying signaling events remain unknown. The conserved heterotrimeric protein phosphatase PP2A controls the timing of events in mitosis, and upstream pathways including Greatwall-Ensa regulate PP2A activity. A role for PP2A in CAR regulation has been less clear, although loss of PP2A in yeast causes defects in cytokinesis. Here, we report that Sds23, an inhibitor of PP2A family protein phosphatases, promotes the symmetric division of fission yeast cells through spatial control of cytokinesis. We found that *sds23*Δ cells divide asymmetrically due to misplaced CAR assembly, followed by sliding of the CAR away from its assembly site. These mutant cells exhibit delayed recruitment of putative CAR anchoring proteins including the glucan synthase Bgs1. Our observations likely reflect a broader role for regulation of PP2A in cell polarity and cytokinesis because *sds23*Δ phenotypes were exacerbated when combined with mutations in the fission yeast Ensa homolog, Igo1. These results identify the PP2A regulatory network as a critical component in the signaling pathways coordinating cytokinesis.

## Introduction

Organisms from yeast to humans build a contractile actomyosin ring (CAR) that drives cytokinesis. The fission yeast CAR assembles from a series of 50-75 precursor structures termed “nodes” in the plasma membrane (Chang *et al.*, 1996; Sohrmann *et al.*, 1996; Wu *et al*., 2006; Almonacid *et al.*, 2009, 2011). Similar precursors have been observed in animal cell CAR assembly (Piekny and Glotzer, 2008). Nodes contain actin- and myosin-regulatory proteins, as well as membrane-binding proteins with the capacity to tether the CAR to the cell cortex (Wu *et al.*, 2006). During mitosis, nodes coalesce into an intact CAR. Past studies have defined the recruitment timing, functional contribution, and spatial organization of CAR proteins in fission yeast (Pollard and Wu, 2010; Laplante *et al.*, 2016; McDonald *et al.*, 2017). However, many questions remain open regarding the upstream signaling pathways that ensure assembly of this complex structure at the correct time and place.

Fission yeast cells proceed through cytokinesis in three steps. The first step is CAR formation, when precursor nodes congress to form an intact ring (Wu *et al.*, 2006). Nodes are positioned in the cell middle by positive signals from the nucleus, and by inhibitory signals from the protein kinase Pom1 at cell tips (Bähler and Pringle, 1998; Daga and Chang, 2005; Celton-Morizur *et al.*, 2006; Padte *et al*., 2006). Both signals converge on the anillin-like scaffold protein Mid1 (Celton-Morizur *et al.*, 2006; Padte *et al.*, 2006; Almonacid *et al.*, 2009; Almonacid *et al.*, 2011). The second step is CAR maturation, when additional proteins are recruited to the division site (Wu *et al.*, 2003; Wu *et al.*, 2006; Pollard and Wu, 2010). During maturation, the CAR must be anchored in place to prevent sliding towards cell ends. Several proteins have been implicated in anchoring the CAR: the cell wall synthase Bgs1 (Arasada and Pollard, 2014; G. Cortés *et al.*, 2015; Sethi *et al*., 2016), the membrane-binding protein Cdc15 (Arasada and Pollard, 2014; McDonald *et al.*, 2015; Snider *et al*., 2017), regulators of the lipid PI(4,5)P_2_ (Snider *et al.*, 2017; Snider *et al.*, 2018), and the scaffolding protein Pxl1 (G. Cortés *et al*., 2015). The third step is CAR constriction, which must be coordinated with septum assembly (Liu *et al.*, 1999; Sipiczki and Bozsik, 2000; Liu *et al.*, 2002). Both CAR constriction and septation contribute to the force required for cytokinesis and cell separation (Jochová *et al.*, 1991; Proctor *et al.*, 2012).

In fission yeast and other systems, protein kinases provide spatial and temporal regulation of cytokinesis. Protein phosphatases, which counter the activity of protein kinases, therefore represent potential regulators of cytokinetic events. In human cells, phosphatases have recently been linked to cytokinesis (Burgess *et al.*, 2010; Cundell *et al.*, 2013). In fission yeast, the phosphatases calcineurin and Clp1 localize to the division site and regulate the timing of cytokinetic events (Trautmann *et al.*, 2001; Mishra *et al.*, 2004; Clifford *et al.*, 2008; Martín-García *et al.*, 2018). Fission yeast mutants in the conserved heterotrimeric phosphatase PP2A have suggested a role for this phosphatase in cytokinesis, but how regulation of PP2A might connect to specific steps in cytokinesis is unclear (Kinoshita *et al.*, 1996; Jiang and Hallberg, 2001; Le Goff *et al.*, 2001; Lahoz *et al.*, 2010; Goyal and Simanis, 2012). PP2A mutants exhibit defects in diverse cellular processes, making it challenging to obtain a clear connection with cytokinesis. PP2A is regulated by a number of upstream factors, which might control specific aspects of PP2A function. We reasoned that mutating these regulators could provide a way to study how this conserved phosphatase regulates cytokinesis without affecting other cellular functions.

In this study, we have identified and characterized cytokinesis defects in cells lacking Sds23, which binds to and inhibits PP2A-related phosphatases (Hanyu *et al.*, 2009; Deng *et al.*, 2017). We found that the CAR assembles off-center in *sds23*Δ mutants, and once assembled slides away from the cell middle due to delayed recruitment of potential anchors. The phenotypes of *sds23*Δ cells are exacerbated when combined with deletion of a second inhibitor of PP2A, Igo1 (Chica *et al.*, 2016), suggesting that Sds23 and Igo1 function together in regulating PP2A-controlled cellular processes.

## Results and Discussion

### Cells lacking Sds23 divide asymmetrically

While studying Sds23 in nutrient signaling (Deng *et al.*, 2017), we observed that *sds23*Δ cells divide with off-center septa (Figure 1A). We quantified this defect by measuring the Cell Half Ratio (Figure 1B, Supplemental Figure 1A) (Snider *et al.*, 2017) and confirmed a significant defect in positioning the division plane for *sds23*Δ cells compared to wild type cells (Figure 1C). We obtained similar results whether Cell Half Ratio was measured by cell length, cross-sectional area, or cell perimeter (Supplemental Figure 1, B and C). At division, *sds23*Δ cells are longer than wild type cells, so we considered the possibility that increased cell length caused the measured division asymmetry. To address this idea, we used a *cdc25-degron-DAmP* (hereafter, *cdc25-dd*) strain, which divides at a similar length to *sds23*Δ (Figure 1A). We found that *cdc25-dd* cells divide as symmetrically as wild type cells, indicating that asymmetric division of *sds23*Δ cells is not due to cell length (Figure 1C). The *sds23*Δ and *cdc25-dd* mutations had additive defects in cell length, but *sds23*Δ *cdc25-dd* double mutant cells displayed the same Cell Half Ratio as *sds23*Δ single mutants (Figure 1C). We conclude that Sds23 regulates the position of the division plane independent of cell size.

**Figure 1.**
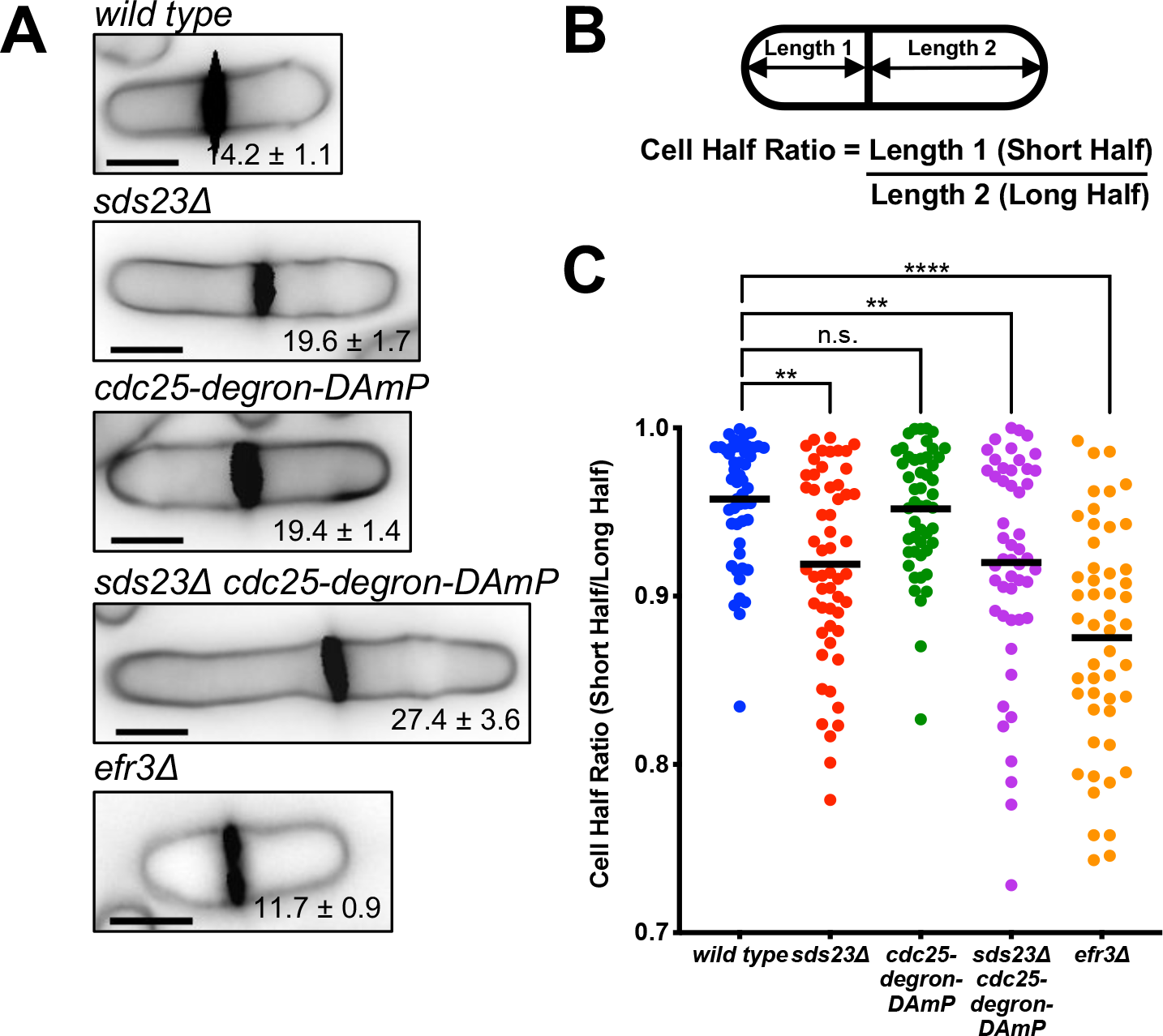
*sds23*Δ cells divide asymmetrically. (A) Representative images of cells of the indicated strains with cell wall stain. Inset is cell length ± standard deviation. Scale bar 5 μm. (B) Schematic depicting calculation of Cell Half Ratio. (C) Cell Half Ratios of the indicated strains as a measure of division asymmetry. ** indicates p-value < 0.01, **** indicates p-value < 0.0001.

### sds23 mutants fail to assemble and to anchor the CAR in the cell middle

We next investigated the cause of misplaced division planes in *sds23*Δ cells. We monitored the position and timing of CAR assembly using *rlc1-mNeonGreen* (hereafter, *rlc1-mNG*) *sad1-mEGFP* cells. Rlc1 marks the precursor nodes and CAR, and Sad1 marks the spindle pole bodies (SPBs), which provide a clock for cytokinetic events (Wu *et al.*, 2003). CAR formation occurred with the same timing in *sds23*Δ and wild type cells, but was misplaced in *sds23*Δ cells (Figure 2, A and B). Misplaced ring assembly is not due to delocalized Pom1, which remained at cell tips in *sds23*Δ cells (Supplemental Figure 1D), or changes in the localization of Cdr2 nodes, which are precursors to cytokinetic nodes (Supplemental Figure 1E). Thus, the off-center CAR phenotype of *sds23*Δ cells does not reflect changes to Pom1-dependent negative spatial cues.

**Figure 2.**
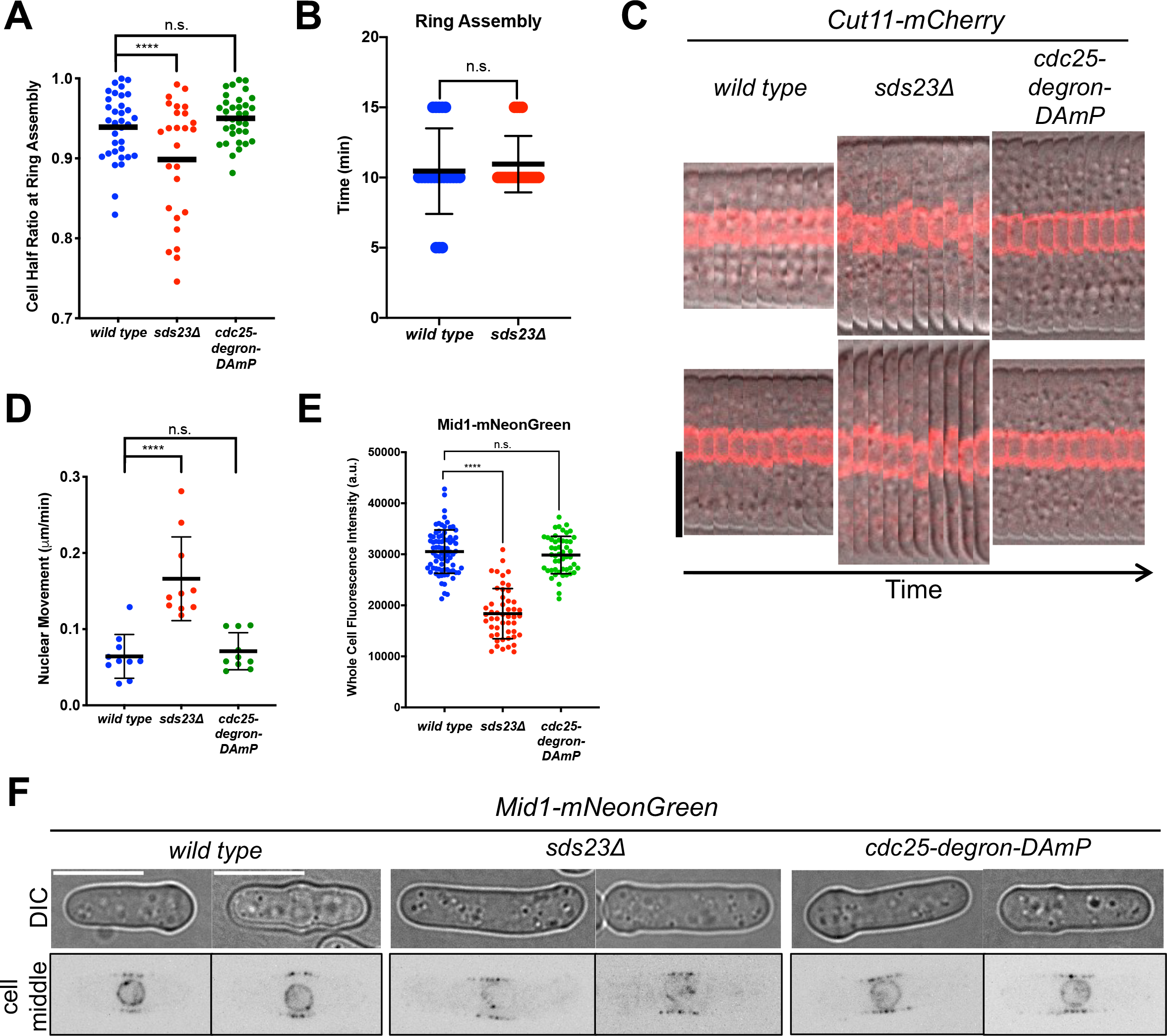
Defects in nuclear positioning and ring assembly in *sds23*Δ cells. (A) Cell Half Ratio at time of ring assembly for the indicated strains. **** indicates p-value < 0.0001. (B) Duration of CAR assembly in wild type versus *sds23*Δ cells. (C) Montages displaying two representative cells of the indicated strains expressing Cut11-mCherry. Cut11-mCherry fluorescence signal has been overlaid onto DIC images to visualize the cell tips. Scale bar 8 μm, images acquired every 3 minutes. (D) Quantification of nuclear movement as a function of time for the indicated strains. **** indicates p-value < 0.0001. (E) Quantification of Mid1-mNeonGreen whole cell fluorescence in wild type, *sds23Δ*, and *cdc25-dd* cells. **** indicates p-value < 0.0001. (F) DIC and middle focal plane inverted fluorescence images of two representative wild type, *sds23Δ*, and *cdc25-dd* cells expressing Mid1-mNeonGreen. Scale bar 8 μm.

In addition to Pom1-dependent negative spatial cues, CAR formation also depends on positive spatial cues from the nucleus. Using Cut11-mCherry to mark the nucleus, we performed time-lapse imaging of wild type and *sds23*Δ cells. In wild type cells, the nucleus remained stationary at the center of cells (Figure 2, C and D), consistent with proper medial placement of the CAR in these cells. In *sds23*Δ cells, the nucleus was not stationary at the cell middle, and instead wandered off-center (Figure 2, C and D). Importantly, nuclear positioning and nuclear movement in elongated *cdc25-dd* cells resembled wild type (Figure 2, C and D). Similar results were obtained by visualizing the localization of the SPB over time (Figure 2E). Microtubules (MTs) connected to the nucleus push against the cell ends to position nuclei in the cell middle (Tran et al., 2001; Daga et al., 2006). To determine if defects in MTs were responsible for the increased nuclear movement, we imaged *sds23*Δ cells with fluorescently tagged MTs. We found that MTs in *sds23*Δ cells have increased dwell times at the cell tips when compared to both wild type and cdc25-dd cells. (Figure 2F). This difference likely contributes to nuclear wandering and off-center CAR formation through mislocalization of positive spatial cues.

At the molecular level, this positive spatial cue is provided by Mid1, which shuttles between the nucleus and precursor nodes to coordinate spatial cues from these two structures (Bähler et al., 1998a; Daga and Chang, 2005; Almonacid et al., 2009). Mid1-mNeonGreen displayed reduced levels in the nucleus and the CAR (Figure 2G and Supplemental Figure 2K), although node localization of Mid1 was not obviously impaired (Figure 2G). Strikingly, the cellular protein level of Mid1 was reduced in *sds23*Δ cells, as measured by both whole cell fluorescence and western blot (Figure 2H and Supplemental Figure 1F). Thus, *sds23*Δ cells have defects in nuclear positioning and in Mid1 regulation. These defects likely lead to misplaced CAR assembly away from the cell middle.

Once assembled, the CAR must be anchored in place during the maturation phase of cytokinesis. In certain mutants, CARs assemble properly but then slide away from the cell middle, resulting in asymmetric division. This defect has been observed in mutants that alter plasma membrane PI(4,5)P_2_ regulation (Snider *et al.*, 2017; Snider *et al*., 2018), or upon mutation of Cdc15, an essential F-BAR domain protein that might tether the CAR to the plasma membrane (Arasada and Pollard, 2014; McDonald *et al.*, 2015; Snider *et al*., 2017). To determine if ring sliding defects occur in *sds23*Δ cells, we performed time-lapse imaging on *rlc1-mNG sad1-mEGFP* cells. After ring formation, CARs in wild type or elongated *cdc25-dd* cells did not move from their initial position. In contrast, CARs slid away from their initial position in *sds23*Δ (Figure 3A). We measured ring displacement (Figure 3B) and found that CARs slid significantly farther in *sds23*Δ cells than in wild type or *cdc25-dd* cells (Figure 3C). In these experiments, CARs slid during the time period corresponding to ring maturation, which occurs after assembly but before constriction. We measured a significantly longer maturation phase in *sds23*Δ versus wild type cells (Figure 3D), indicating that this CAR stage requires Sds23 to proceed normally.

We next tested if this Sds23 function in cell division symmetry connects with PI(4,5)P_2_ regulation by combining *sds23*Δ with *efr3*Δ. Efr3 is a scaffold for PI-4 kinase, and *efr3*Δ cells were recently shown to exhibit ring sliding (Snider *et al.*, 2017) (Figure 1, A and C). The *sds23*Δ *efr3*Δ double mutant did not show additive defects in division asymmetry (Figure 3E), but this result was difficult to interpret because *sds23*Δ *efr3*Δ double mutant cells lysed at a high frequency (Figure 3F). This synthetic fitness defect suggests that Sds23 may function through a different pathway from PI(4,5)P2 regulation.

**Figure 3.**
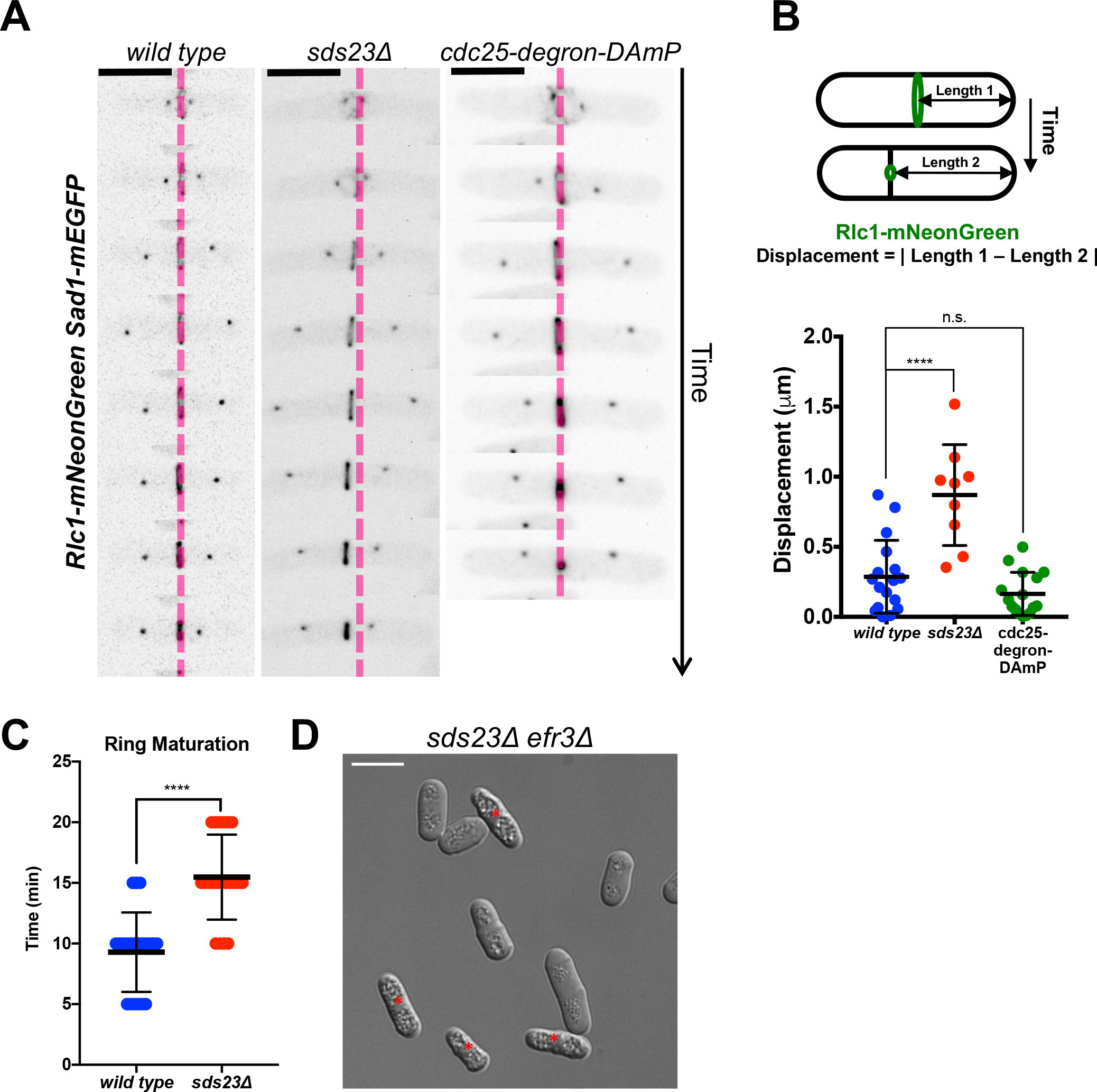
CAR sliding in *sds23*Δ cells. (A) Timelapse montages displaying a representative wild type, *sds23*Δ, and *cdc25-degron-DAmP* cell expressing Rlc1-mNeonGreen and Sad1-mEGFP. The first image of each montage is the time of SPB splitting. The pink dotted line indicates initial ring position for all montages. Images are inverted fluorescence of sum projections. Scale bar 8 μm. (B) Top: Schematic depicting ring displacement measurement. Bottom: Quantification of cytokinetic ring displacement. **** indicates p-value < 0.0001. (C) Duration of CAR maturation in wild type versus *sds23*Δ cells. **** indicates p-value < 0.0001. DIC image of *sds23Δ efr3Δ*. Red asterisks indicate dead cells.

### Sds23 is required for timely recruitment of CAR anchors

Reduced recruitment of proteins that anchor the CAR in place could explain both the ring sliding and extended maturation phase of *sds23*Δ mutant cells. To test this hypothesis, we measured the levels of 22 cytokinesis proteins at the CAR using live-cell microscopy of wild type or *sds23*Δ cells (Supplemental Figure 2, A-V). Several proteins that might contribute to ring anchoring were present in reduced levels at the CAR of *sds23*Δ cells. These proteins include Bgs1, a transmembrane cell wall enzyme, and Imp2 and Rga7, which both contain membrane-binding F-BAR domains (Supplemental Figure 2, A, B and P) (Demeter and Sazer, 1998; Martin-Garcia *et al.*, 2014; Arasada and Pollard, 2015; McDonald *et al.*, 2016). We did not detect altered levels of Cdc15, another F-BAR domain containing protein that is implicated in ring anchoring (Supplemental Figure 2C) (Roberts-Galbraith *et al.*, 2009; McDonald *et al.*, 2015).

Past work has suggested that delayed recruitment of Bgs1, which synthesizes 1,3-beta-glucan for the primary septum, can lead to ring sliding (Arasada and Pollard, 2014). We measured the timing of GFP-Bgs1 recruitment in *sds23*Δ and wild type cells using the SPB clock (Wu *et al.*, 2003). Bgs1 recruitment to the division site was delayed 10-15 minutes in *sds23*Δ compared to wild type cells (Figure 4, A and B), consistent with a similar extension in the maturation phase of cytokinesis in this mutant. We observed similar delays in Imp2-GFP (~5 minutes) (Figure 4, C and D) and Rga7-GFP (~10-15 minutes) (Figure 4, E and F) recruitment in *sds23*Δ cells compared to wild type cells. Such delays were specific to these factors, as other proteins such as Rlc1 were recruited to the CAR with similar kinetics in *sds23*Δ and wild type cells (Supplemental Figure 1, G and H). We conclude that Sds23 is required for the timely recruitment of specific putative CAR anchoring proteins to prevent ring sliding.

**Figure 4.**
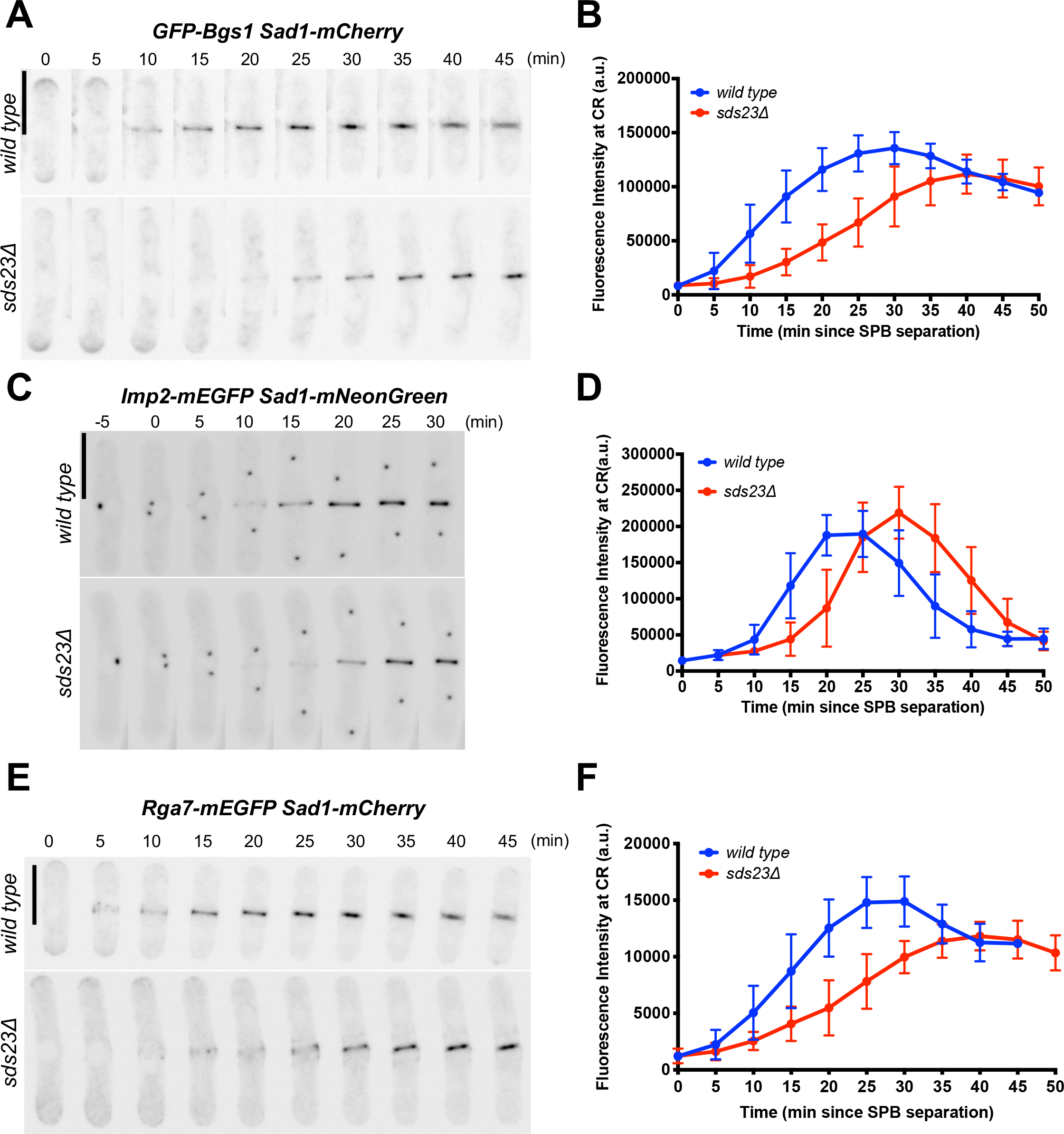
Delayed recruitment of CAR anchors in *sds23*Δ cells. (A) Timelapse images of GFP-Bgs1 recruitment in wild type versus *sds23*Δ cells. Images are inverted fluorescence of sum projections. Time “0” indicates SPB splitting as determined by Sad1 signal. Note, a 561 nm laser was used to image Sad1-mCherry, but only the signal from the 488 nm laser is displayed. Scale bar 8 μm. (B) Quantification of GFP-Bgs1 recruitment during cytokinesis in wild type versus *sds23*Δ cells. (C) Timelapse images for Imp2-mEGFP recruitment, as in panel A. (D) Quantification of Imp2-mEGFP recruitment during cytokinesis in wild type versus *sds23*Δ cells. Timelapse images for Rga7-mEGFP recruitment, as in panel A. (F) Quantification of Rga7-mEGFP recruitment during cytokinesis in wild type versus *sds23*Δ cells.

### Genetic interactions between Sds23 and the Greatwall Kinase Pathway

Sds23 physically associates with PP2A and PP6 phosphatases, and can inhibit PP6 phosphatase activity (Hanyu *et al.*, 2009; Deng *et al.*, 2017). Similar to other organisms, the fission yeast endosulfine protein Igo1 inhibits PP2A activity (Chica *et al.*, 2016). The Greatwall kinase Ppk18 phosphorylates and activates Igo1, raising the possibility that Ppk18-Igo1 and Sds23 might provide dual regulatory arms for PP2A phosphatases (Figure 5A) (Chica *et al.*, 2016). Both *igo1*Δ and *ppk18*Δ mutants alone did not display defects in cell shape, cell length at division, or division symmetry under normal growing conditions, consistent with previous reports (Figure 5, B-E) (Chica *et al.*, 2016). However, combining either mutation with *sds23*Δ led to synthetic defects. First, the percentage of cells with aberrant morphology, specifically bent and curved cells, was increased in the *sds23*Δ *igo1*Δ and *sds23*Δ *ppk18*Δ double mutant cells compared to single mutants (Figure 5C). Second, the double mutants divided at a larger cell size than the single mutants (Figure 5D), consistent with both pathways acting on PP2A to control mitotic entry. Lastly, the Cell Half Ratio was even further reduced for double mutant cells, but we note that the double mutants were not significantly different from *sds23*Δ alone despite this trend (Figure 5E). Taken together, these results suggest that Sds23 and Igo1 share functions related to cell morphology and cell size, but Sds23 plays a more prominent role in cytokinesis. Consistent with this notion, elongating *igo1*Δ by combining with cdc25-dd had no effect on division symmetry (Supplemental Figure 3, A-C) and the *sds23*Δ *igo1*Δ did not have additive defects in nuclear movement (Supplemental Figure 3D). Synthetic defects between Sds23 and Ppk18-Igo1 support a broad role for PP2A-family regulators in cell polarity, cell size, and division plane positioning.

**Figure 5.**
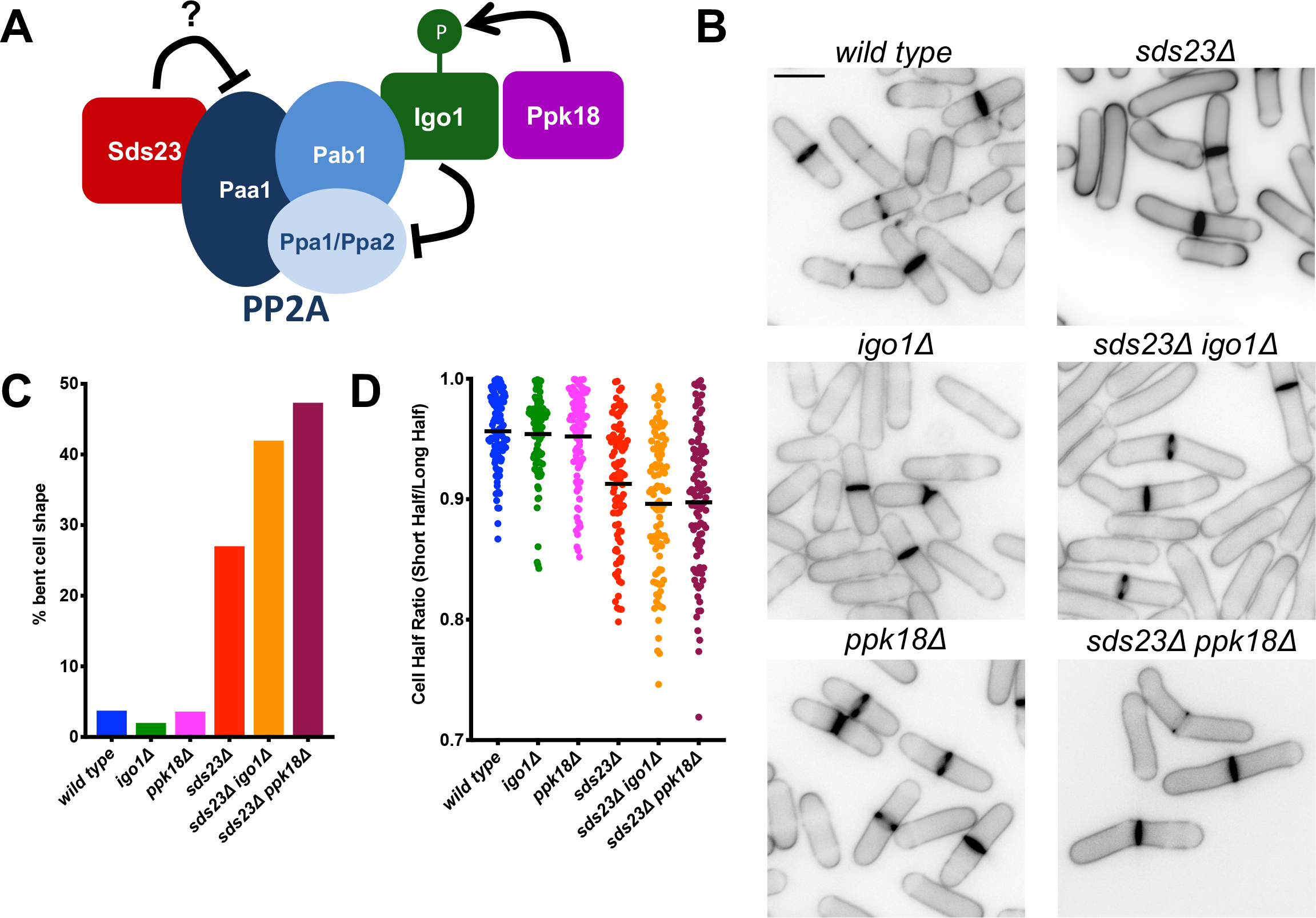
Genetic interactions between Sds23 and Ppk18-Igo1. (A) Schematic depicting Igo1 and Sds23 interactions with Protein Phosphatase 2A (PP2A). Phosphorylation of Igo1 by Ppk18 promotes interaction with PP2A (Hanyu *et al.*, 2009; Chica *et al.*, 2016). (B) Representative images of cells of the indicated strains with cell wall stain. Scale bar 8 μm. (C) Quantification of bent cell shape phenotype for the indicated strains. (D) Cell Half Ratios of the indicated strains.

### Conclusions

Our results define a new role for regulation of PP2A in cytokinesis and cell division. Past work has shown that *sds23*Δ cells exhibit increased levels of PP2A activity (Hanyu *et al.*, 2009), and our results show that these cells have defects in regulating key cytokinesis proteins such as Mid1 and Bgs1. These defects likely underlie misplaced assembly of the CAR and its subsequent sliding away from the cell middle. Future experiments will address if these cytokinesis proteins are direct targets of dysregulated PP2A or if their dysregulation occurs downstream of PP2A itself. PP2A has been proposed to counter the SIN in fission yeast (Jiang and Hallberg, 2000; Jiang and Hallberg, 2001; Le Goff *et al.*, 2001; Lahoz *et al.*, 2010), therefore, Sds23 and Igo1 action on PP2A may alter the balance of PP2A versus SIN activity within the cell, leading to the division phenotypes observed in this study. We note that reduced PP2A activity in *ypa2*Δ has been shown to suppress some SIN mutants, but *ypa2*Δ did not correct the Cell Half Ratio of *sds23*Δ cells (Supplemental Figure 3E). In fact, *ypa2*Δ cells alone exhibited a Cell Half Ratio defect, showing that strict regulation of PP2A activity is critical for division plane positioning.

The ability of an intact CAR to slide laterally within the plane of the plasma membrane indicates a remarkable level of structural integrity and stability, even though many components are exhibiting rapid flux as shown by photobleaching experiments (Pelham and Chang, 2002). It seems likely that the CAR is anchored in place by multiple tethers, including the components implicated in this study as well as previous work (Arasada and Pollard, 2014; McDonald *et al.*, 2015; Snider *et al.*, 2017). Such a multi-component anchoring system may facilitate a spatial control system that is robust during dynamic processes including septum synthesis, vesicle delivery, and CAR constriction.

Our work places this PP2A regulatory network in the growing context of protein phosphatases that control cytokinesis. Given the roles of PP2A in diverse cellular functions, it will be interesting to know how potential cytokinesis targets are regulated with specificity in time and space. The partially overlapping roles of Sds23 and Igo1, as shown by synthetic genetic defects, raise possibilities for dynamic control of cytokinesis by PP2A during cell cycle progression. The Greatwall-endosulfine pathway including Ppk18-Igo1 is best known for regulation of mitotic entry (Mochida *et al.*, 2010; Burgess *et al.*, 2010; Chica *et al.*, 2016), and Sds23 has also been linked to cell cycle progression through phenotypes such as cell size (Ishii *et al*., 1996; Jang *et al*., 1997; Deng *et al*., 2017). These functional connections to the cell cycle suggest that timely regulation of PP2A by these proteins could ensure proper cytokinesis. Both Sds23 and Igo1 are known to be hyperphosphorylated, which may serve to regulate their interactions with PP2A (Hanyu *et al.*, 2009; Chica *et al.*, 2016). We also note that the Ppk18-Igo1 pathway is upregulated during nitrogen stress (Chica *et al.*, 2016), and Sds23 functions in pathways upregulated during glucose depletion and energy stress (Yakura *et al.*, 2006; Hanyu *et al.*, 2009; Jang *et al.*, 2013; Saitoh *et al.*, 2015; Deng *et al.*, 2017). Thus, it will be interesting to determine how these pathways contribute to cytokinesis when cells encounter environmental stresses linked to their functions. Given the conservation of PP2A complexes and functions in yeast through humans, phosphatase regulation by proteins such as those described here might contribute to robust cytokinesis in diverse cell types and organisms.

## Materials and Methods

### Yeast Strains and Growth

Standard *S. pombe* media and methods were used (Moreno *et al.*, 1991). Strains used in this study are listed in Supplemental Table 1. Gene tagging and deletion were performed using PCR and homologous recombination (Bähler et al., 1998b), and integrations were verified by colony PCR, microscopy, and/or western blot. To cross the sterile mutant *sds23*Δ, a pJK148 plasmid with *sds23*+ or *sds23-mNeonGreen*+ was integrated at the *leu1* locus of a *sds23*Δ*::natR leu1-32* strain, and *leu1-32 natR* progeny or *natR* progeny lacking *mNeonGreen* signal were selected from subsequent crosses.

### Microscopy

All image analysis was performed on ImageJ (National Institutes of Health) (Schneider *et al*., 2012). For Figure 1A and Supplemental Figure 3A, cells were grown at 32°C, stained with Blankophor (MP Biomedicals), mounted on a glass slide under a coverslip, and middle focal plane images were obtained at room temperature. Imaging was performed on a Deltavision Imaging System (Applied Precision/GE Healthcare), which used a customized Olympus IX-71 inverted wide-field microscope, a Photometrics Cool-SNAP HQ2 camera, Insight solid-state Illumination unit and 1.42 NA Plan Apo 60x oil objective. For Figure 3D, cells were grown at 32°C, mounted on a glass slide under a coverslip, and middle focal plane images were obtained at room temperature using the Deltavision Imaging System. For Supplemental Figure 1A, cells were grown at 32°C, mounted on a glass slide under a coverslip, and middle focal plane images were acquired at room temperature using the Deltavision Imaging System. For Supplemental Figure 1B, cells were grown at 32°C, mounted on a glass slide under a coverslip, and z-sectioning was performed with images acquired every 0.5 μm for 6 μm (12 steps). Images were acquired using the Deltavision imaging system and z-stacks were processed by iterative deconvolution in SoftWorx Software (Applied Precision/GE Healthcare). To calculate FWHM, a rectangular ROI of 160 pixels × 48 pixels was drawn around cells that were between 15-17 μm in length and fluorescence profile for Cdr2-mEGFP was plotted. A Gaussian distribution was fitted to this fluorescence profile for each cell using the Least Squares Regression method in Graphpad Prism, and FWHM was calculated by the following equation: 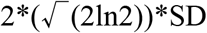, where SD is the standard deviation. Each point represents a single cell and statistical analysis was performed by unpaired t-test with Welch’s Correction. For Figure 5B, cells were grown at 32°C, stained with Blankophor (MP Biomedicals), mounted on a glass slide under a coverslip, and middle focal plane images were obtained using a Nikon Eclipse Ti microscope equipped with an Andor Zyla camera, Perfect Focus and a Tokai Hit stage incubator maintained at 32°C and using a Plan Apo **λ** 1.4 NA aperture 60x oil objective.

For Cell Half Ratio analysis, cell length of each daughter cell was measured from Blankophor stained images, and Cell Half Ratio was calculated by dividing the short half by the long half. For statistical analysis in Figure 1C, a one-way Anova with Tukey’s multiple comparisons test was performed. For Figures 2C, 3A, 4A, 4C, 4E, and Supplemental Figure 1C cells were imaged at 32°C using a Andor W1 Spinning Disk Confocal, set up on a Nikon Eclipse Ti inverted microscope stand with Perfect Focus and Tokai Hit stage incubation system using a Plan Apo **λ** 1.4 NA aperture 60x oil objective. For imaging, ~1 μl of cells were mounted on EMM4S agarose pads (1.4% InCert™ Agarose, Lonza) and sealed under a coverslip with VALAP (Vaseline, Lanolin, Parrafin). For Figures 2B and 3C, ring assembly and maturation timing were determined from time-lapse movies of Rlc1-mNeonGreen and Sad1-mEGFP expressing cells that were imaged every 5 minutes. Ring assembly was determined as the time it took from SPB splitting to form an intact CR. Ring maturation was determined as the time between ring assembly and prior to constriction. Each point represents a single cell. For statistical analysis, a one-way ANOVA with Tukey’s multiple comparisons was used. For Figure 2F, cells were grown at 32°C, mounted on a glass slide under a coverslip, and maximum intensity projections were generated using 0.5-μm focal planes throughout the entire cell (13 steps, 6 μm total). Images were obtained using the Andor W1 Spinning Disk Confocal and a Tokai Hit stage incubator maintained at 32°C and using a Plan Apo λ 1.45 NA 100x oil objective.

For images in Figure 2C, single focal plane images in the cell middle were acquired every 3 minutes. To measure nuclear movement a line was drawn from the center of the nucleus to the cell tip, and the length of this line was measured at every time point. Nuclear movement (μm/min) was quantified as the average difference between those lengths between every time point, divided by 3 minutes. Each point in Figure 2D is the average for a single cell with 10 cells measured per strain. For statistical analysis of nuclear movement, a one-way ANOVA with Tukey’s multiple comparisons test was used. For images in Figure 3A, images were acquired every 5 minutes and sum intensity projections were generated using 0.5-μm focal planes throughout the entire cell (11 steps, 5 μm total). For displacement measurements in Figure 3B, the length of one cell half was measured at ring formation and the length of that same cell half was measured at final ring constriction. The difference between these two values was quantified as the ring displacement (μm) (See Figure 3B schematic). Each dot in Figure 3B corresponds to a single cell and for statistical analysis a one-way ANOVA with Tukey’s multiple comparisons test was used.

For Figures 4A, 4C, 4E, and Supplemental Figure 1C images were acquired every 5 minutes using the Andor W1 Spinning Disk Confocal and sum intensity projections were generated using 0.2-μm focal planes throughout the entire cell (27 steps, 5 μm total) using a Plan Apo λ 1.4 NA 60x oil objective. For Supplemental Figure 2 A-T, cells were imaged in liquid medium under a coverslip, and images were acquired on the Andor W1 Spinning Disk Confocal at 32°C using a Plan Apo λ 1.4 NA 60x oil objective. Sum projection images (non-deconvolved) were used for all quantitative analyses in Figure 4, Supplemental Figure 1D, and Supplemental Figure 2. For Figure 4, Supplemental Figure 1D, and Supplemental Figure 2, an ROI corresponding to the cytokinetic ring was drawn to measure integrated density. To measure background signal, the same ROI was moved to a place with no cells and this background signal was subtracted. For statistical analyses in Supplemental Figure 2, unpaired t-tests with Welch’s correction were performed and p-values were determined by two-tailed distribution. For all statistical analyses, Graphpad Prism was used. For all statistical analyses, a single asterisk (*) denotes p-values < 0.05, two asterisks (**) indicate p-values < 0.01, three asterisks indicate p-value <0.001 and four asterisks (****) indicate p-value < 0.0001.

## Acknowledgements

We thank members of the Moseley lab for comments on the manuscript. We thank the Biomolecular Targeting Core (BioMT) (P20-GM113132) and the Imaging Facility at Dartmouth for use of equipment. We also thank Jian-Qiu Wu, Juan Carlos Ribas, Dan McCollum, and Vladimir Sirotkin for sharing strains. This work was supported by grants from the American Cancer Society (RSG-15-140-01) and the National Institute of General Medical Sciences (R01GM099774) to J.B.M. K.L.S. was supported by a T-32 Training Grant (T32GM008704).

---

**Supplemental Figure 1.**
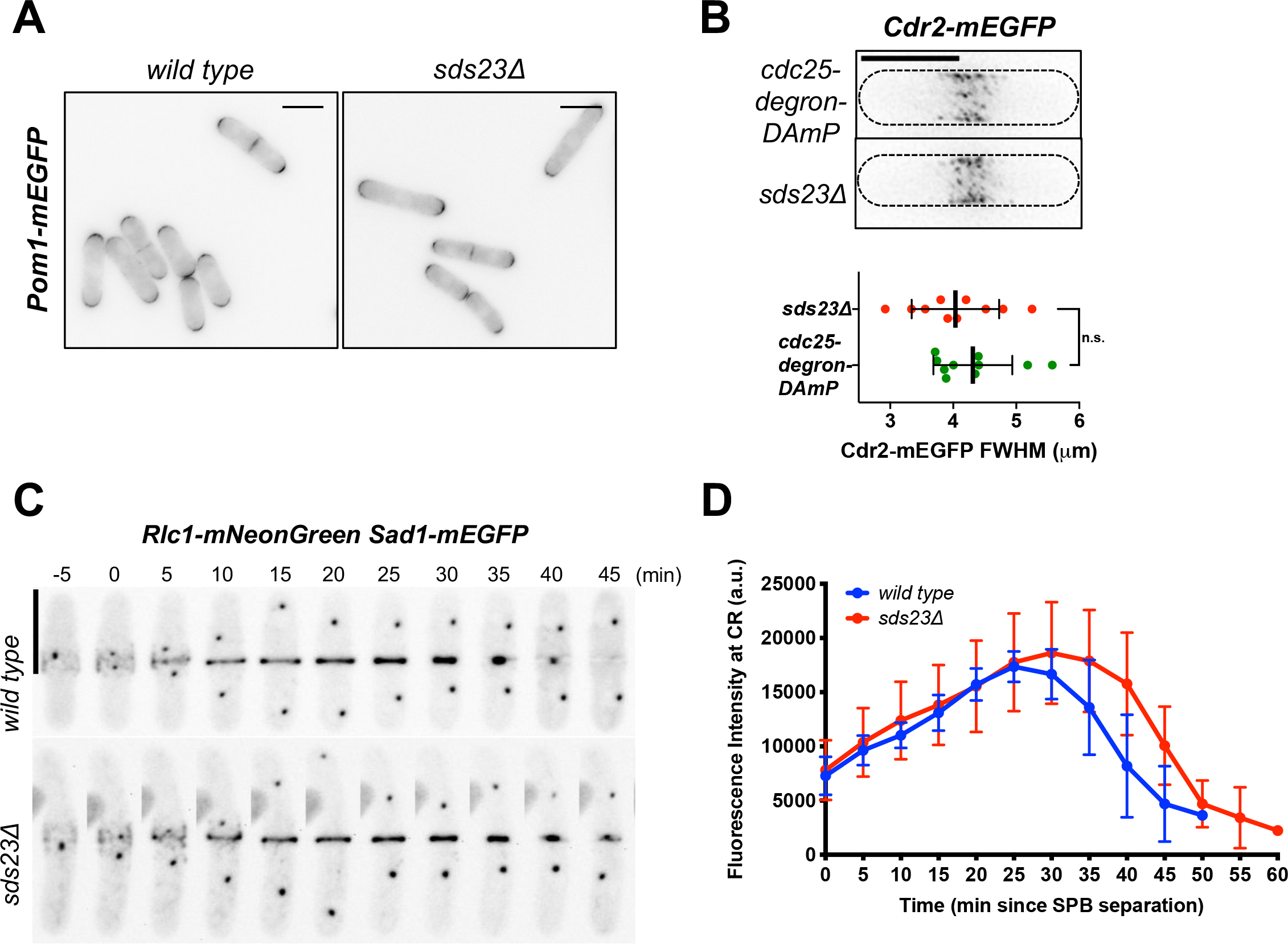
Distribution of nodes and myosin recruitment are unaffected in *sds23*Δ. (A) Inverted fluorescence images of Pom1-mEGFP in wild type and *sds23*Δ cells. Scale bar 5 μm. (B) Top: Inverted max projection images of *cdc25-degron-DAmP* and *sds23*Δ cells expressing Cdr2-mEGFP. Scale bar 8 μm. Bottom: Full-Width Half Maximum (FWHM) of Cdr2-mEGFP signal comparing spread of nodes in *cdc25-degron-DAmP* and *sds23*Δ cells. (C) Time lapse images of wild type and *sds23*Δ cells expressing Rlc1-mNeonGreen and Sad1-mEGFP. Images are inverted fluorescence of sum projections. Scale bar 8 μm. (D) Quantification of Rlc1-mNeonGreen over time for wild type and *sds23*Δ.

**Supplemental Figure 2.**
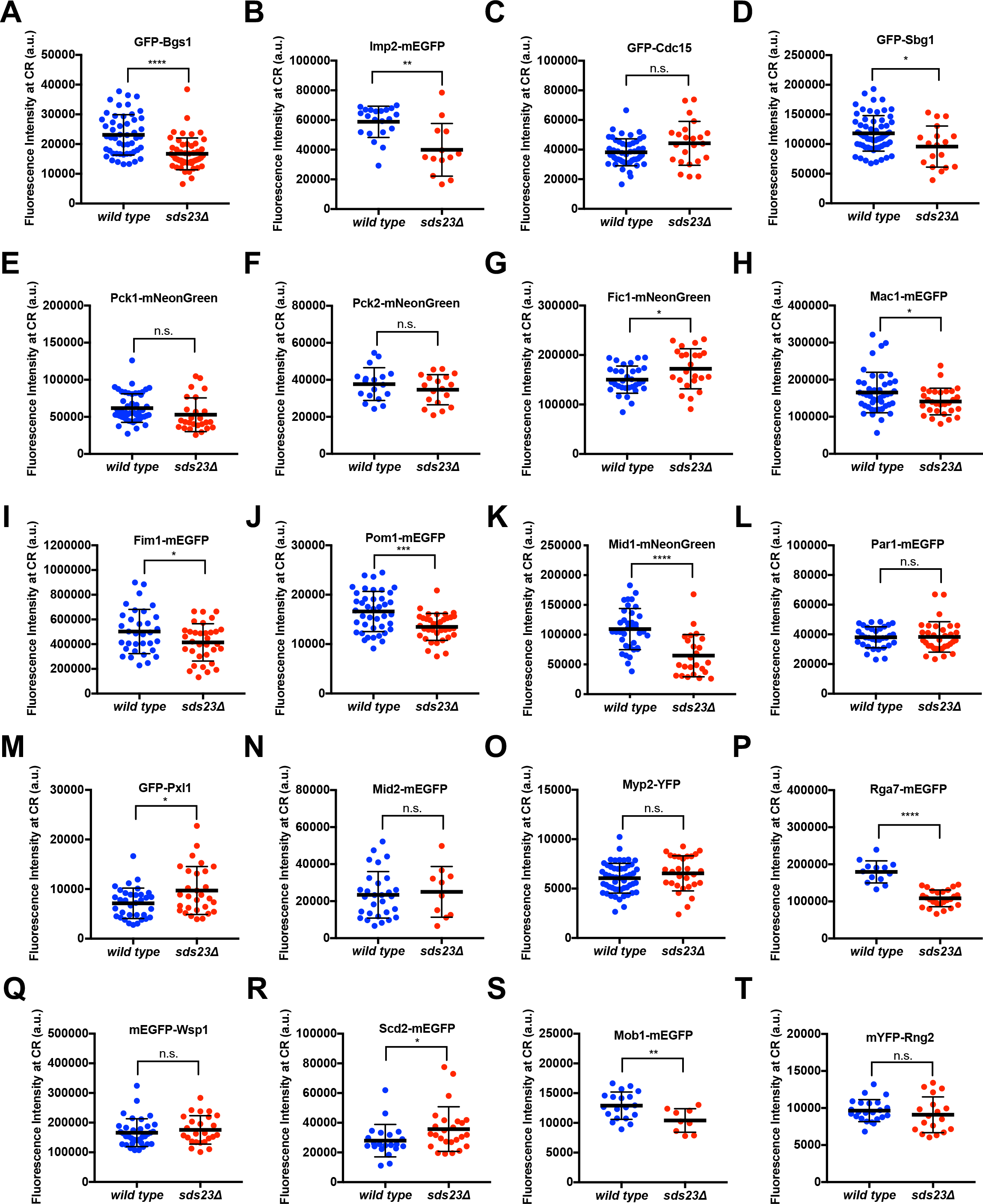
Screen for localization defects for putative CAR anchor proteins. (A-T) Quantification of the indicated CAR proteins in wild type and *sds23*Δ cells. * indicates p-value < 0.05, ** indicates p-value < 0.01, *** indicates p-value < 0.001, and **** indicates p-value < 0.0001.

**Supplemental Figure 3.**
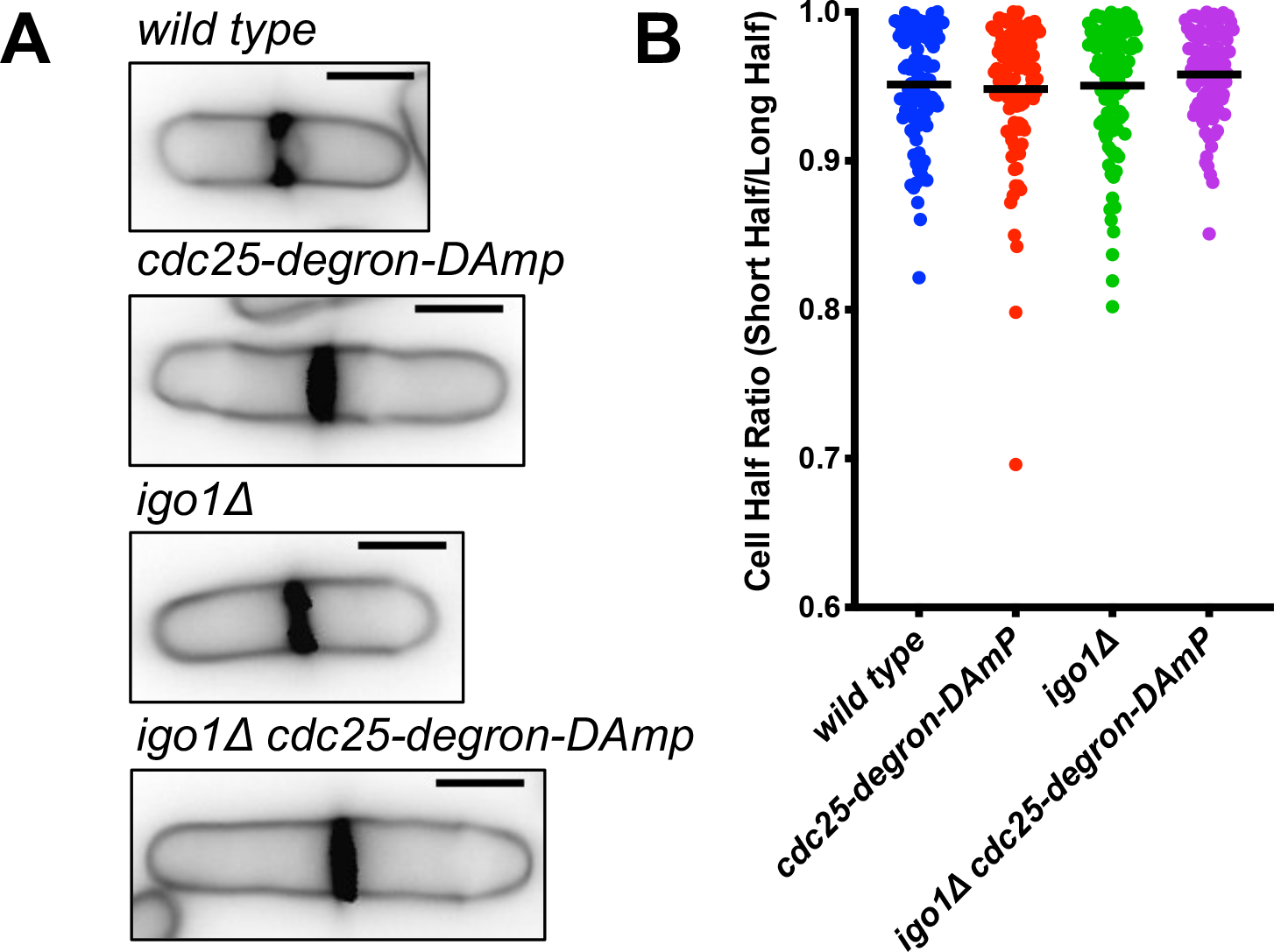
Elongating *igo1*Δ cells does not cause cytokinesis defects. (A) Representative images of cells of the indicated strains with cell wall stain. Scale bar 5 μm. (B) Cell Half Ratios of the indicated strains as a measure of division asymmetry.

**Supplemental Table 1. Strains used in this study.**

